# Punctual ecological changes rather than global factors drive species diversification and the evolution of wing phenotypes in *Morpho* butterflies

**DOI:** 10.1101/2021.04.18.440344

**Authors:** Nicolas Chazot, Patrick Blandin, Vincent Debat, Marianne Elias, Fabien L. Condamine

## Abstract

Assessing the relative importance of geographical and ecological drivers of evolution is paramount to understand the diversification of species and traits at the macroevolutionary scale. Here, we use an integrative approach, combining phylogenetics, biogeography, ecology, and quantified phenotypes to investigate the drivers of both species and phenotypic diversification of the iconic Neotropical butterfly genus *Morpho*. We generated a time-calibrated phylogeny for all known species and inferred historical biogeography. We fitted models of time-dependent (accounting for rate heterogeneity across the phylogeny) and paleoenvironment-dependent diversification (accounting for global effect on the phylogeny). We used geometric morphometrics to assess variation of wing size and shape across the tree, and investigated their dynamics of evolution. We found that the diversification of *Morpho* is best explained when considering multiple independent diversification dynamics across the tree, possibly associated with lineages occupying different microhabitat conditions. First, a shift from understory to canopy was characterized by an increased speciation rate partially coupled with an increasing rate of wing shape evolution. Second, the occupation of dense bamboo thickets accompanying a major host-plant shift from dicotyledons towards monocotyledons was associated with a simultaneous diversification rate shift and an evolutionary “jump” of wing size. Our study points to a diversification pattern driven by punctual ecological changes instead of a global driver or biogeographic history.

## Introduction

Investigating the rates of phenotypic evolution and the relationships between phenotypes and species ecology can shed light on the drivers of time and geographic patterns of diversity. Previous studies have demonstrated that rates of both species and phenotypic diversification vary widely through time and among clades at all taxonomic scales (e.g. Venditti *et al*., 2011; Eastman *et al*., 2011; Rabosky & Adams, 2012; Rabosky *et al*., 2013; Rabosky *et al*., 2014; Cooney & Thomas, 2020). These variations have resulted in the striking heterogeneity in species and phenotypic diversity observed across the tree of life. Such variations may eventually be coupled, indicating an interaction between the processes of species and phenotypic diversifications. Studies investigating such coupling have yielded contrasted results. Some of them support an association between specific and phenotypic diversification (e.g. in salamanders: Rabosky & Adams, 2012; fish: Rabosky *et al*., 2013; vertebrates: Cooney & Thomas, 2020), while others found no support for this relationship (e.g. in lizards: Rabosky *et al*., 2014; squirrels: Zelditch *et al*., 2015; reef fishes: Price *et al*.,2015; snakes: Lee *et al*., 2016). For example in squirrels, Zelditch *et al*. (2015) suggested that species diversification was geographically driven while phenotypic diversification was ecologically driven, resulting in a decoupling of the two dynamics.

A correlation between species and phenotypic diversification rates is notably expected in some specific cases. For example, adaptive radiations – rapid adaptive diversification in a variety of ecological niches – are expected to produce bursts of diversification and phenotypic evolution especially during the initial stages of diversification (Schluter, 2000; Gavrilets & Losos, 2009). Speciation rate increases when a large number of ecological niches are vacant while phenotypes rapidly evolve in response to the diversity of ecological opportunities. Strong correlation between speciation rates and phenotypic diversification may also be found when the focal trait directly drives reproductive isolation. For example, the evolution of male genitalia, involved in mating, may facilitate reproductive isolation between populations (see Langerhans *et al*., 2016 for a review). Correlated dynamics leading to a lower rate of diversification can also be predicted. For example, if extinction probability is biased with respect to phenotype leading to a non-random loss of variation in a particular clade, both species and morphological diversity should show a correlated drop down (Foote, 1997). In this study, we assess the role of multiple ecological causes of variations in rates of species and wing diversification and the extent to which these variations are coupled, by focusing on the case of the butterfly genus *Morpho* (Nymphalidae).

The genus *Morpho* comprises 30 species (Blandin & Purser, 2013), which are amongst the largest butterflies in the Neotropics and are well known for their blue iridescent wing coloration. Several ecological factors have already been suggested as potential drivers of diversification and phenotypic evolution. Previous biogeographic estimations suggested that *Morpho* butterflies originated and started diversifying in the Andes (Penz *et al*., 2012, Blandin & Purser 2013), before spreading across the Neotropics. There is also evidence that *Morpho* lineages separated early in their history into two microhabitats (DeVries *et al*., 2010; Chazot *et al*., 2016). One clade is composed of species that tend to fly high, often above the forest canopy, with some species typically harbouring gliding flight behaviour such as *M. cisseis* and *M. hecuba*. The remaining species mostly fly within the first meters above ground in the understory (DeVries *et al*., 2010; Chazot *et al*., 2016). Finally, according to Cassildé *et al*. (2010) and Penz *et al*. (2012), the genus *Morpho* was ancestrally feeding on monocotyledons, and two major host-plant shifts occurred during its diversification: after the first divergence event, one of the two clades shifted to dicotyledon host-plants and, within this clade, a subclade subsequently reversed to the monocotyledons.

Here we focus on the wings of *Morpho*, which are at the crossroad of multiple selective pressures and tightly linked to species diversification. Typically, wing colour patterns can be involved in camouflage, aposematism or courting behaviours (Naisbit *et al*., 2001; Merrill *et al*., 2011). Wings also allow flight, enabling dispersal, foraging, predator escape, mating or host-plant searching (Dudley, 2002). Hence butterfly wings are under strong natural and/or sexual selection and may be associated with variations of speciation rate (Ortiz-Acevedo *et al*., 2020). Both size and shape are important aspects of wing morphology. They both strongly affect the performance of flight behaviours (Dudley, 2002; Le Roy *et al*.,2019) and therefore might be closely associated to habitat use, dispersal strategies or host-plant searching. Besides, fore and hind wings can be functionally differentiated, for example during flight (Grodnitsky *et al*., 1994; Le Roy *et al*., 2020), which may lead to uncorrelated patterns of diversification.

To investigate whether species and phenotypic diversification dynamics are coupled and to identify potential drivers of variations, we inferred a time-calibrated molecular phylogeny of the genus that we combined to a dataset of geographical distributions and morphometric measurements of wing size and shape. We applied an integrative approach and addressed the following questions: (1) Have rates of phenotypic diversification varied across the tree? We investigated potential variations in rate of phenotypic diversification among clades using phenograms and models of trait evolution to compare evolutionary rates for wing size and shape. (2) Is species diversification better explained by global processes or clade-specific (ecological) factors? First, we fitted different models of species diversification testing for global drivers of diversification, specifically past temperatures and Andean orogeny. Second, we compared these global drivers to models in which species diversification varied according to clade-specific ecological factors (microhabitat and major shifts of host-plants) and/or variations of phenotypic diversification identified in the first step. (3) Can we explain the variations in diversification rates by historical biogeography? We performed ancestral areas estimation in order to assess whether variations in phenotypic evolutionary rates or species diversification rates may be associated with specific biogeographic events.

## Material and methods

### Time-calibrated phylogeny

Phylogenetic relationships and divergence time were inferred with Bayesian inference. We concatenated DNA data for one mitochondrial (COI) and four nuclear genes (CAD, EF-1α, GAPDH and MDH) using published sequences (Cassildé *et al*., 2012; Penz *et al*., 2012; Chazot *et al*., 2016) retrieved from GenBank, generating a molecular dataset of a total length of 5001 nucleotides. Our dataset includes all *Morpho* species (i.e. 30 species *sensu* Blandin, 2007). *Morpho helenor*, which harbours many subspecies, is distributed throughout the entire Neotropical region, resulting in unresolved biogeographic reconstructions in preliminary analyses. To help resolving the biogeographic inferences, *M. helenor* was represented in the biogeographic analyses by six subspecies that each occupies a distinct Neotropical area. For all other analyses, we pruned all subspecies of *M. helenor* but one in order to keep a single branch for the species. We also included 11 outgroups to root and calibrate the tree (see Supporting Information S1) on the basis of the most comprehensive nymphalid phylogeny to date (Wahlberg *et al*., 2009).

To simultaneously estimate the topology and branching times of the phylogeny we used a Bayesian relaxed-clock approach as implemented in *BEAST* 1.8.2 (Drummond *et al*., 2012). To choose the best partitioning strategy and the corresponding substitution models, we ran *PartitionFinder* 1.1.1 (Lanfear *et al*., 2012) allowing all possible partitions and models implemented in *BEAST*. Three subsets were defined: the first included position 1 and 2 of all genes and followed a GTR+I+Γ model, the second included position 3 of all nuclear fragments and followed a GTR+Γ model, and the third including the position 3 of the mitochondrial fragment and followed a TrN+Γ model. We implemented an uncorrelated lognormal relaxed clock model. Given the lack of *Morpho* fossil we relied on secondary calibrations to calibrate the molecular clock. Penz *et al*. (2012) calibrated the divergence between *Morpho* and its sister groups using a unique calibration point from Wahlberg *et al*. (2009), and a normal distribution for the corresponding prior. However, Sauquet *et al*. (2012) showed that using a single secondary calibration prior could yield biased estimates. Hence, we used a set of seven calibrations defined by uniform priors bounded by the 95% credibility intervals (95% CI) estimated by Wahlberg *et al*. (2009) (see Supporting Information S1). We implemented a Yule process for the tree prior, and we ran the phylogenetic analyses for 30 million Markov chain Monte Carlo (MCMC) generations. We checked for chain convergence using *Tracer* 1.6, as indicated by effective sample size (ESS) values. Finally, we used *TreeAnnotator* 1.8.2 (Drummond *et al*., 2012) to select the maximum clade credibility (MCC) tree with median age values calculated from the posterior distribution of branch lengths, applying a 20% burn-in.

### Morphological data

Our morphological dataset (published by Chazot *et al*. 2016) consists in the size and shape of the fore and hind wings, as assessed by morphometric measurements. A total of 911 collection specimens of both sexes and representing all *Morpho* species were photographed. Wing shape was described using landmarks and semi-landmarks placed at vein intersections and wing margins, respectively (see Chazot *et al*., 2016 for details), which were superimposed with *tpsRelw* (Rohlf, 1993). Wing size was measured using the log-transformed mean centroid size per species. Importantly, for analyses involving wing shape we used the residuals of a multivariate regression of species mean Procrustes coordinates on species mean centroid size (log-transformed), which allows focusing on the non-allometric shape variation. Similar analyses were performed separately on the fore and hind wings. All analyses were performed on males and females separately. As we found divergent patterns among sexes, we show the results for males and females separately. No female *M. niepelti* was available. This species was therefore pruned from the tree for all analyses involving female data.

### Dynamics of phenotypic diversification

We investigated whether the evolutionary rates of wing size and shape have varied among subclades across the phylogeny.

#### Wing size

We first visualized the evolution of traits through time using the *phenogram* function in *PHYTOOLS* 0.5-20 (Revell, 2012), which represents the trait values inferred at each node along a time axis. Second, we investigated the dynamics of wing size evolution across lineages using the method implemented in the function *rjmcmc.bm* available in *GEIGER* 2.0.6 (Harmon *et al*., 2008; Eastman *et al*., 2011) for univariate traits. This method uses Bayesian analyses and reversible-jump MCMC to infer the number and the location of shifts of morphological diversification dynamics. We fitted and compared three different models of trait evolution: (1) a single-rate Brownian model (BM); (2) a relaxed model of Brownian evolution in which a trait evolved according to distinct Brownian-motion models across the tree (rBM); and (3); a model in which trait evolution can also occur at punctual “jumps”, i.e. brief periods of rapid evolution at any branch in the phylogeny (jBM). We ran models on both the MCC and a posterior distribution of trees. For the MCC tree analysis, we ran for each model one MCMC of 30 million generations, sampling every 3,000 generations. We checked for convergence of each run using *CODA (*Plummer *et al*. 2020), and computed the ESS. We applied a 25% burn-in and compared the three models using the Akaike’s Information Criterion for MCMC samples *aicm* and *aicw* implemented in *GEIGER* (Supporting Information S2). To assess the robustness of the inferences to branch length uncertainties, we repeated the analysis on a posterior distribution of trees and summarized the results. We sampled 100 trees with a topology identical to that of the MCC tree from the posterior distribution. For each tree, we ran the three models but reduced the MCMC to 10 million generations, calculated the *aicm* score, the mean *aicm* pairwise differences between models and the position of rate shifts and jumps. We summarized the results by calculating the frequency of shifts and jumps at nodes across the posterior distribution. These results are hereafter referred to as shift/jump posterior tree frequencies.

#### Wing shape

Some authors have used the scores on the first PC-axis as a univariate shape measure (e.g. Rabosky *et al*., 2014; Thacker, 2014) to investigate shifts in evolutionary rates for multidimensional traits, but this may lead to spurious results (Uyeda *et al*., 2014). We rather investigated variations in rates of shape evolution across the phylogeny in a multivariate way using the function *compare.evol.rate* from *GEOMORPH* (Adams, 2014; Denton & Adams, 2015). It allows testing whether species assigned to different ecological factors have significantly different rates of shape evolution, by comparing the ratio between the rates of each group to a null distribution of ratios obtained through simulation of a unique neutral evolutionary rate (two or more factors can be tested). When more than two factors are included, the function performs a global test for the significance of the multiple rates model compared to a one-rate model, but also assesses the significance of differences among each pair of factors. We used this factor assignment to define monophyletic subgroups with potentially divergent evolutionary rate from the background rate. We first tested all models with one shift, i.e. all species belonging to one subclade (each subclade had a minimum of three species) are assigned to one group, and the rest of the species assigned to another group. If two or more subclades were identified as having a rate of evolution significantly different from that of the background, we identified the subclade with the highest ratio (hence the greatest shift). Then we ran again *compare.evol.rate* on all possible combinations of two shifting subclades that include the first identified shift. A two-shift model was considered significant if at least the two shifting subclades showed a significant difference with the background rate when considering the pairwise comparisons. Given the relatively small size of our phylogeny, we limited our analysis to two shifts (Supporting Information S3-S4). As for wing size, we repeated the analysis on both the MCC tree and a posterior distribution of trees with identical topologies. We summarized the results from the posterior distribution by calculating the frequency of significant shifts at nodes across the trees, and refer to these as posterior tree frequencies.

### Dynamics of species diversification

We compared two types of species diversification models: (1) diversification rates varying according to global factors, i.e. factors virtually affecting all lineages, and (2) diversification rates varying at specific clades characterized by clade-specific ecological factors. For each type we investigated different factors (see below). All models were compared using their AIC scores to identify the model that best explains the diversification of the genus *Morpho*.

#### Global drivers of diversification

We tested the role of temperature fluctuations and of the paleo-elevation of the Andes on species diversification by using birth-death models that allow speciation and extinction rates to vary according to a past environmental variable itself varying through time (Condamine *et al*., 2013). For each paleoenvironmental variable, we designed three models to be tested: *(i)* the speciation rate varies exponentially with the environment and the extinction rate is constant, *(ii)* the speciation rate is constant and the extinction rate varies exponentially with the environment, and *(iii)* both speciation and extinction rates vary exponentially with the environment. We repeated these three models with a linear dependence to the environmental variable, instead of exponential dependence. For temperature we relied on the well-known Cenozoic temperature dataset published by Zachos *et al*. (2008). The orogeny of the Andes is a highly complex process, with important differences in uplift tempo and mode from the south of Central Andes to Northern Andes (Blandin & Purser, 2013, and references therein). Several general phases have been identified from the late Eocene to present, but they are difficult to synthetize in a unique model. As Blandin & Purser (2013) suggested that the early diversification of the *Morpho* occurred along the proto-Central Andes, we used the model of surface uplift inferred by Leier *et al*. (2013) for the eastern cordillera of the southern Central Andes to test the possible influence of Andean orogeny on the diversification of the *Morpho*. We used the R-package *PSPLINE* 1.0-17 to reconstruct smooth lines of the paleo-data for each environmental variable. The smooth line is introduced in the birth-death model to represent the variation of the environment through time. Given the dated phylogeny, the model then estimates speciation and extinction rates, as well as their respective variations according to the environment (Condamine *et al*., 2013). These analyses were performed on 200 trees randomly sampled from the posterior distribution generated by BEAST.

#### Clade-specific drivers of diversification

We assessed whether the diversification rates across the genus *Morpho* have varied among specific clades using models of time-dependent diversification. To do so we used the method developed by Morlon *et al*. (2011), which allows partitioning diversification rates into independent dynamics (a backbone and different subclades). We compared different partitioning schemes according to three events: (1) the microhabitat change (from understory to canopy), (2) the shift of wing shape evolutionary rate, and (3) the reverse shift to monocotyledon host-plants (also identified as a punctual evolutionary jump of wing size at the stem). Because the evolutionary rate shift of wing shape is nested within the microhabitat shift (see Results), we could not test both combined. Instead, each of those shifts was combined to the monocotyledon host-plant shift with a two-shift model of diversification rate. For each subclade and the remaining backbone, we fitted the following models: *(i)* constant speciation rate and no extinction, *(ii)* time-dependent speciation rate and no extinction, *(iii)* constant speciation and extinction rates, *(iv)* time-dependent speciation rate and constant extinction rate, *(v)* constant speciation rate and time-dependent extinction rate, and *(vi)* time-dependent speciation and extinction rates. Time dependency was modelled using an exponential function of time. The stem branch of each subclade was included in the subclades and excluded from the backbones but we kept the node of the divergence (speciation event) of the subclade within the backbones. The root of the tree was excluded from the analyses. The analysis was performed on the MCC tree, since partitioning the tree requires defining clades *a priori*, which entails a fixed topology.

### Historical biogeography

To assess where and when diversification occurred, we estimated ancestral areas using the dispersal-extinction-cladogenesis (DEC, Ree and Smith, 2008) model as implemented in the R-package *BioGeoBEARS* 0.2.1 (Matzke, 2014). The analyses were performed using the MCC tree (outgroups removed) and included six subspecies of *M. helenor* (each subspecies was assigned to its current distribution).

The distribution of *Morpho* is restricted to South America and Central America (all Neotropics except the Caribbean Islands). A geographic model was incorporated to include operational areas, defined as geographic ranges shared by at least two or more species and delimited by geological, oceanic or landscape features, which may have acted as barriers to dispersal. The model comprised 7 component areas: (A) Central America, (B) trans-Andean South-America, (C) slopes of northern Andes, (D) eastern slopes of central Andes, Orinoco-Amazonian basin north of the Amazon, including the Guyanas, (E) Amazonian basin, south of the Amazon River, and (F) Atlantic forest.

An adjacency matrix was designed whilst taking into account the geological history and the biological plausibility of combined ranges (Supporting Information S6). Distributional data were compiled from monographies (Blandin, 2007). We excluded distribution margins overlapping with adjacent areas. For example, *M. marcus* and *M. eugenia* are mainly found in lowlands but their distributions reach the Andean slopes up to altitudes of 700-800m. Nevertheless, we did not consider these as species occupying the Andean biogeographic areas. By contrast, a species such as *M. sulkowskyi*, which occurs between 1500 and 3500m high in the Andes was considered as an Andean species. We also set a maximum of 3 areas per node to be constitutive of an ancestral range. We fitted two different DEC models, one that assumed equal dispersal probabilities among all areas and one that included time-stratified matrices of varying dispersal probabilities (Supporting Information S6). We compared the likelihoods of both reconstructions to select the model best explaining the current pattern of species distribution.

## Results

### Divergence times

We estimated that the genus *Morpho* diverged (stem age) from its sister genus *Caerois* 38.05 Ma (95% CI=35.48-39.20 Ma) and the first event (crown age) of diversification was recovered at 28.12 Ma (95% CI=25.22-31.24 Ma; Supporting Information S1). These divergence time estimates are slightly older than those estimated by Penz *et al*. (2012) and Chazot et al. (2019) who found an average divergence from *Caerois* around 32.00 Ma and 29.08 Ma respectively. This difference probably results from prior choices for calibrating the trees (see Material and Methods).

### Dynamics of phenotypic diversification

#### Wing size

Both analyses on the MCC and posterior distribution of trees found similar results. We found no support for any shift in rate of wing size diversification. However, we found support for an evolutionary jump. For females the model jBM was highly supported for both wings, with a highly probable (posterior tree frequency [PF] of 0.99, Supporting Information S2) evolutionary jump at the root of the clade including the species *M. absoloni*, *M. aurora*, *M. zephyritis*, *M. rhodopteron*, *M. sulkowskyi*, *M. lympharis*,*M. aega*, and *M. portis* (subclade portis). Phenograms show that in this subclade, female wings are on average 34% smaller than in the other *Morpho* species for both fore and hind wings (Fig. 1). This is all the more striking as the sister clade (including *M. amathonte*, *M. menelaus*, and *M. godartii*) contains some of the largest species of the genus (e.g. *M. amathonte* has a wingspan of 10–15 cm). For males, the *portis* clade exhibits the same trend, but the support for the evolutionary jump is lower than for females (PF_forewing_=0.76, PF_hindwing_=0.71, respectively, Supporting Information S2). Males wings in the portis clade were on average 30 and 32% smaller for fore and hind-wing respectively.

**Figure 1.**
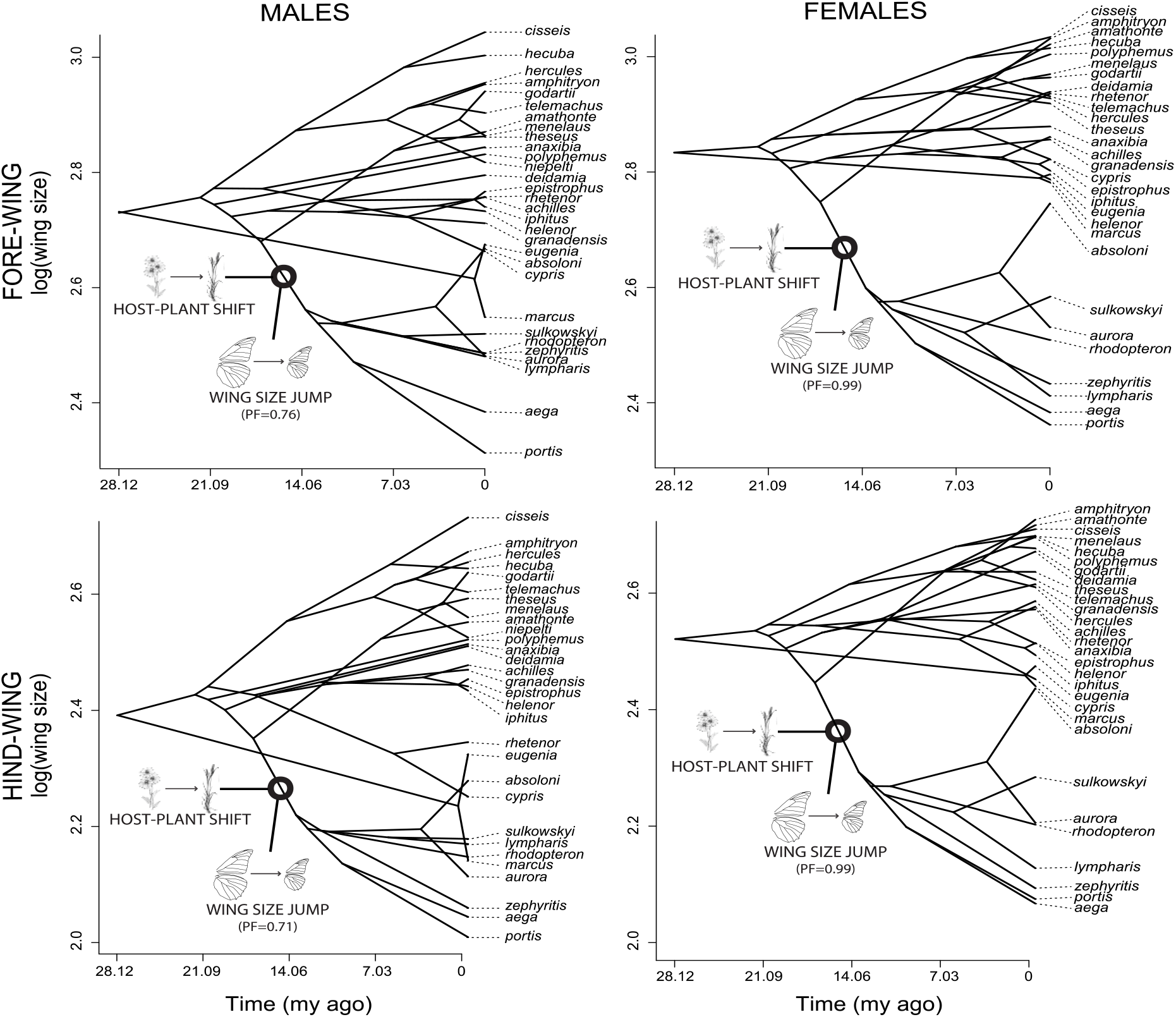
Phenograms for wing size (log scale) for males (left panels) and females (right panels). The top panels are the forewings, and the bottom panels are the hindwings. Wing size values are reconstructed at the nodes and plotted on a time scale. Phylogenetic relationships are projected into the phenogram. The position (branch) where the main host-plant shift and significant wing size jump happened is also shown. PF values indicate the frequency at which each jump was found across the posterior distribution of trees.

#### Wing shape

We found support for two shifts of evolutionary rate for male hindwing, in both cases towards lower rate of evolution. These subclades encompass *M. helenor*, *M. achilles*, and *M. granadensis* (Fig. 2, Supporting Information S3) on one side, and *M. godartii*, *M. menelaus*, and *M. amathonte* on the other. This result was supported by the analyses with the MCC tree. The analyses performed on the posterior tree distribution found a moderate support for these shifts, with PF of 0.62 and 0.77, respectively. For females and for both wings the subclade encompassing *M. theseus*, *M. amphitryon*, *M. telemachus* and *M. hercules* exhibited the greatest shift (highest ratio) (Fig. 2, Supporting Information S4). This shift corresponds to a large increase in rate of evolution (forewing ratio=181.74, hindwing ratio=184.49 in the MCC analysis), i.e. wing shape evolving faster within this group than the other *Morpho*. This result was strongly supported by posterior distribution analyses, with PF of 0.96 and 0.99 for fore and hind-wing, respectively.

**Figure 2.**
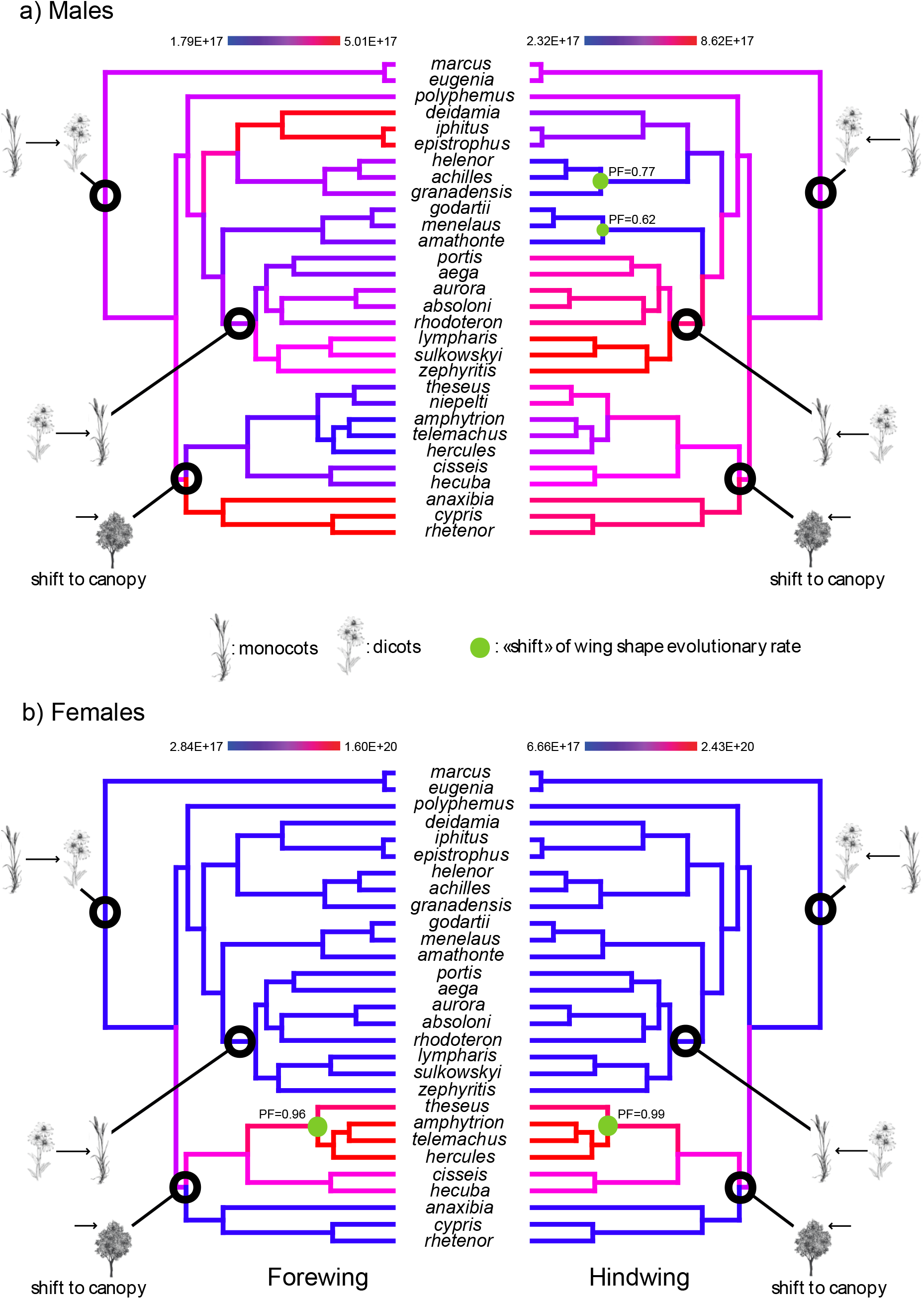
Rate of wing shape diversification for a) males and b) females. Branches of the phylogenies are coloured according to the evolutionary rate inferred at the nodes using the R package GEOMORPH. Green points indicate the changes in the rate of wing shape evolution and black points the evolutionary jumps of wing size. Only shifts with a posterior tree frequency higher than 0.5 are shown. PF values indicate the frequency at which each shift was found across the posterior distribution of trees. On these phylogenies some major evolutionary events including important host-plant shifts and microhabitat shifts are also indicated.

### Dynamics of species diversification

#### Global drivers of diversification

In the best model accommodating for Central Andean paleo-altitudes, speciation rates were negatively dependent on the paleo-altitude and extinction rates were constant (Table 2a). This model leads to a continuous decrease in speciation rate towards the present, suggesting that *Morpho* diversification was high during the early stages of the orogeny but the rise in altitude did not lead to any increased opportunities for speciation over time. We also found a significant correlation between *Morpho* diversification and temperature compared to a null model (Table 2b). The best fitting paleoclimatic model indicates that speciation rate was positively correlated with temperature variation while extinction remained constant. This means that speciation rate was high during the initial stages of diversification when the temperatures were warmer but globally decreased during the last 14 million years as the Earth cooled down (Zachos *et al*., 2008).

**Table 1.** Summary results obtained from fitting three models of trait evolution on 100 trees, using the *rjmcmc.bm* function as implemented in the R package GEIGER on a) males and b) females. *bm*=single Brownian rate, *rbm*=relaxed Brownian rates, *jbm*=jumps of Brownian rates. AICbm, AICrbm, AICjbm = mean AIC score across the 100 trees for all three models. ΔAIC bm-rbm, ΔAIC bm-jbm, ΔAIC rbm-jbm = pairwise AIC differences between models for each tree.

**a) Males**
| | AICbm | AICrbm | AICjbm | ΔAIC bm-rbm | ΔAIC bm-jbm | ΔAIC rbm-jbm |
| --- | --- | --- | --- | --- | --- | --- |
| Forewing | -23.92 (18.72) | -27.78 (10.33) | 13.41 (26.32) | 3.85 (21.70) | -37.34 (34.54) | -41.20 (30.02) |
| Hindwing | -14.43 (20.28) | -18.62 (14.58) | 14.94 (24.28) | 4.18 (24.02) | -29.37 (29.76) | -33.56 (28.21) |

**b) Females**
| | AICbm | AICrbm | AICjbm | ΔAIC bm-rbm | ΔAIC bm-jbm | ΔAIC rbm-jbm |
| --- | --- | --- | --- | --- | --- | --- |
| Forewing | -22.36 (10.44) | -17.74 (9.40) | -34.70 (1.60) | -4.61 (15.73) | 12.33 (10.98) | 16.95 (9.56) |
| Hindwing | -15.44 (17.06) | -13.30 (22.83) | -32.80 (1.17) | -2.14 (29.62) | 17.36 (16.88) | 19.50 (23.01) |

**Table 2.** Paleoenvironmental-dependent diversification analyses using paleoaltitude (a) and Cenozoic temperature (b) data. Mean parameter and standard error estimates are presented for each model. Best-fitting model, as determined via a combination of the lowest AIC and ΔAIC (see main text) highlighted in bold. In our best-fit paleoaltitude-dependent model, speciation is negatively correlated to Andean orogeny over time (adding extinction as a parameter did not improve the model fit). Likewise, speciation is positively correlated to temperature variation over time (allowing extinction to vary with temperature did not improve the likelihood). *λ* = speciation rate (in events/Myr/lineage); μ = extinction rate (in events/Myr/lineage); α = rate of variation of the speciation according to the relevant paleoenvironmental variable; β = rate of variation of the extinction according to the paleoenvironmental variable; NP = number of parameters in each model.

**a) Paleoaltitude models**
| Models | Dependency | NP | logL | AIC | ΔAIC | $\lambda$ | $\alpha$ | $\mu$ | $\beta$ |
| --- | --- | --- | --- | --- | --- | --- | --- | --- | --- |
| $\lambda$ Alti. and no $\mu$ | Linear | 2 | -97.50<br>$\pm 0.085$ | 199.00<br>$\pm 0.171$ | 0.00 | 0.190<br>$\pm 0.004$ | -3.60E-05<br>$\pm 1.39$E-06 | - | - |
| $\lambda$ Alti. and no $\mu$ | Exponential | 2 | -97.68<br>$\pm 0.083$ | 199.37<br>$\pm 0.166$ | 0.37 | 0.210<br>$\pm 0.004$ | -3.06E-04<br>$\pm 5.76$E-06 | - | - |
| $\lambda$ Alti. and $\mu$ constant. | Linear | 3 | -97.49<br>$\pm 0.085$ | 200.99<br>$\pm 0.170$ | 1.99 | 0.190<br>$\pm 0.0003$ | -3.47E-05<br>$\pm 8.75$E-07 | 5.15E-04<br>$\pm 4.17$E-04 | - |
| $\lambda$ Alti. and $\mu$ constant | Exponential | 3 | -97.69<br>$\pm 0.083$ | 201.37<br>$\pm 0.166$ | 2.37 | 0.220<br>$\pm 0.004$ | -3.11E-04<br>$\pm 5.68$E-06 | 9.09E-07<br>$\pm 7.91$E-07 | - |
| $\lambda$ constant and $\mu$ Alti | Exponential | 3 | -97.75<br>$\pm 0.104$ | 201.51<br>$\pm 0.207$ | 2.51 | 0.080<br>$\pm 3.16$E-04 | - | 42038.92<br>$\pm 11873.33$ | -7.48E-03<br>$\pm 2.62$E-04 |
| $\lambda$ Alti. and $\mu$ Alti | Exponential | 4 | -96.94<br>$\pm 0.094$ | 201.87<br>$\pm 0.187$ | 2.87 | 0.880<br>$\pm 0.160$ | -5.00E-04<br>$\pm 2.00$E-05 | 6.820<br>$\pm 1.159$ | -1.89E-03<br>$\pm 4.72$E-05 |
| $\lambda$ Alti. and $\mu$ Alti. | Linear | 4 | -97.42<br>$\pm 0.086$ | 202.83<br>$\pm 0.171$ | 3.84 | 0.210<br>$\pm 0.004$ | -3.83E-05<br>$\pm 1.18$E-06 | 0.070<br>$\pm 0.007$ | -2.38E-05<br>$\pm 2.32$E-06 |
| $\lambda$ constant and $\mu$ Alti. | Linear | 3 | -98.56<br>$\pm 0.099$ | 203.12<br>$\pm 0.198$ | 4.12 | 0.080<br>$\pm 0.001$ | - | 0.030<br>$\pm 0.005$ | -1.14E-05<br>$\pm 1.76$E-06 |

b) Paleoclimate models
| Models | Dependency | NP | logL | AIC | $\Delta AIC$ | $\lambda$ | $\alpha$ | $\mu$ | $\beta$ |
| --- | --- | --- | --- | --- | --- | --- | --- | --- | --- |
| $\lambda$ Temp. and no $\mu$ | Exponential | 2 | -96.65<br>$\pm 0.095$ | 197.30<br>$\pm 0.189$ | 0.00 | 0.030<br>$\pm 4.13E-04$ | 0.179<br>$\pm 1.91E-03$ | - | - |
| $\lambda$ Temp. and no $\mu$ | Linear | 2 | -96.72<br>$\pm 0.097$ | 197.44<br>$\pm 0.195$ | 0.13 | 0.015<br>$\pm 5.50E-04$ | 0.013<br>$\pm 1.17E-04$ | - | - |
| $\lambda$ Temp. and $\mu$ constant | Exponential | 3 | -96.58<br>$\pm 0.092$ | 199.15<br>$\pm 0.184$ | 1.85 | 0.030<br>$\pm 3.93E-04$ | 0.210<br>$\pm 3.62E-03$ | 0.043<br>$\pm 0.004$ | - |
| $\lambda$ Temp. and $\mu$ constant | Linear | 3 | -96.67<br>$\pm 0.097$ | 199.33<br>$\pm 0.195$ | 2.03 | 0.023<br>$\pm 1.48E-03$ | 0.021<br>$\pm 1.34E-03$ | 0.046<br>$\pm 0.007$ | - |
| $\lambda$ Temp. and $\mu$ Temp. | Linear | 4 | -96.42<br>$\pm 0.095$ | 200.85<br>$\pm 0.189$ | 3.54 | 0.017<br>$\pm 1.11E-03$ | 0.027<br>$\pm 8.07E-04$ | 0.290<br>$\pm 0.012$ | -0.037<br>$\pm 0.002$ |
| $\lambda$ Temp. and $\mu$ Temp. | Exponential | 4 | -96.55<br>$\pm 0.092$ | 201.10<br>$\pm 0.183$ | 3.80 | 0.032<br>$\pm 7.99E-04$ | 0.205<br>$\pm 4.57E-03$ | 0.156<br>$\pm 0.040$ | -0.164<br>$\pm 0.019$ |
| $\lambda$ constant and $\mu$ Temp. | Exponential | 3 | -98.57<br>$\pm 0.103$ | 203.14<br>$\pm 0.205$ | 5.84 | 0.081<br>$\pm 2.96E-04$ | - | 4.47E-08<br>$\pm 3.08E-09$ | 0.004<br>$\pm 2.19E-04$ |
| $\lambda$ constant and $\mu$ Temp. | Linear | 3 | -98.57<br>$\pm 0.103$ | 203.14<br>$\pm 0.205$ | 5.84 | 0.081<br>$\pm 2.97E-04$ | - | 1.09E-04<br>$\pm 1.08E-04$ | -1.37E-05<br>$\pm 1.37E-05$ |

#### Clade-specific dynamics of diversification

The best-partitioned models included a shift of diversification rate for the host-plant shift and for the canopy shift (Table 3, Supporting Information S5). Under this configuration, the diversification of the clade that shifted to monocotyledon host-plants was best modelled by a speciation rate decreasing through time combined with no extinction, and the diversification of the canopy clade was best modelled by a constant speciation rate with no extinction (Table 3, Fig. 3). For the remaining backbone lineages the best fitting model was a time-dependent speciation and extinction. The resulting net diversification rate (speciation minus extinction) of this backbone was high during the very early stages of diversification but rapidly decreased through time and became negative *ca*. 25 Ma, implying a declining diversity (Fig. 3). Around 22 Ma, the net diversification rate became positive again and reached zero at the present. This model of partitioned dynamics of diversification outperformed any model involving a global driver of diversification. Indeed, the multi-rate time-dependent model better fit the diversification of *Morpho* (AICc=191.69) than the temperature-dependent model (AICc=197.3, ΔAIC=5.61) and the altitude-dependent model (AICc=199.0, ΔAIC=7.31).

**Table 3.** Results of model comparison for the five time-dependent diversification analyses presented, with mean parameter estimates for each model. *λ* = speciation rate (in events/Myr/lineage); α = parameter of rate variation for speciation; μ = extinction rate (in events/Myr/lineage); β = parameter of rate variation for extinction; NP = number of parameters in each model; AICc = corrected Akaike information criterion; logL = log-likelihood.

| Clade partition | Models | NP | logL | AIC | $\lambda$ | $\alpha$ | $\mu$ | $\beta$ | Joint logL | Joint AIC |
| --- | --- | --- | --- | --- | --- | --- | --- | --- | --- | --- |
| background | BVAR<br>DVAR | 4 | -35.68 | 79.36 | 0.063 | 0.237 | 0.079 | 0.228 |  |  |
| monocots | BVAR | 2 | -21.78 | 47.55 | 0.014 | 0.213 | - | - | -88.84 | 191.69 |
| canopy | BCST | 1 | -31.39 | 64.77 | 0.083 | - | - | - |  |  |
| background | BCST | 1 | -50.91 | 103.83 | 0.072 | - | - | - |  |  |
| monocots | BVAR | 2 | -21.78 | 47.55 | 0.014 | 0.213 | - | - | -92.83 | 193.66 |
| shape shift | BCST | 1 | -20.14 | 42.28 | 0.095 | - | - | - |  |  |
| background | BCST | 1 | -73.75 | 149.49 | 0.081 | - | - | - | -95.52 | 197.05 |
| monocots | BVAR | 2 | -21.78 | 47.55 | 0.014 | 0.213 | - | - |  |  |
| whole | BCST | 1 | -98.40 | 198.81 | 0.081 | - | - | - | -98.40 | 198.81 |
| background | BCST | 1 | -78.18 | 158.36 | 0.078 | - | - | - | -98.32 | 200.63 |
| shape shift | BCST | 1 | -20.14 | 42.28 | 0.095 | - | - | - |  |  |
| background | BCST | 1 | -67.01 | 136.03 | 0.080 | - | - | - | -98.40 | 200.80 |
| canopy | BCST | 1 | -31.39 | 64.77 | 0.083 | - | - | - |  |  |

**Figure 3.**
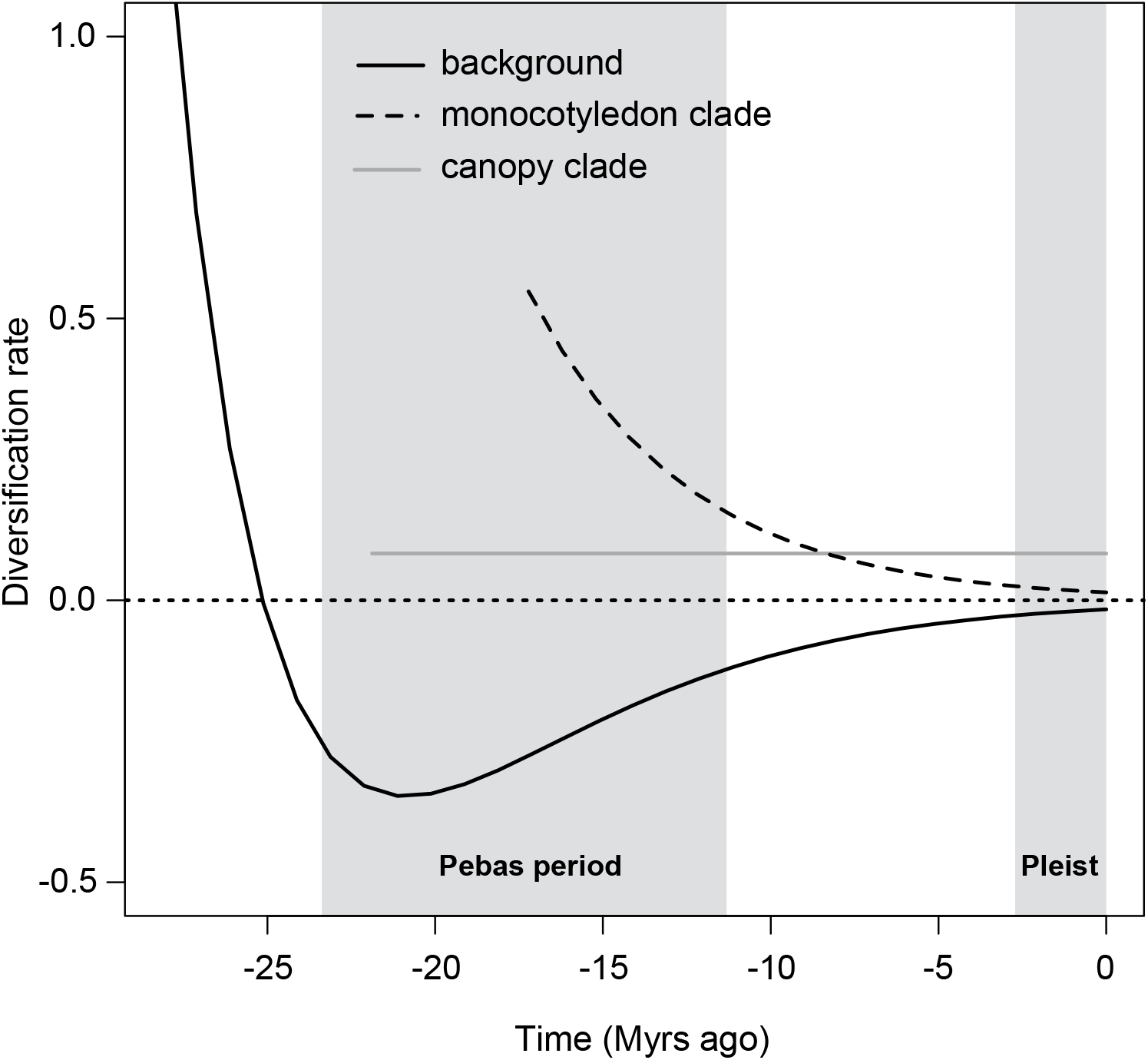
Estimation of the temporal dynamics of diversification for the genus *Morpho*. Diversification rates (speciation minus extinction) for the best models identified for the different subclades (canopy and monocotyledon) and the remaining lineages (background). The early background diversification is elevated and decreases through time until it becomes negative in the early Miocene. The canopy clade has constant rates of diversification, while the monocotyledon clade conforms to an early-burst pattern with high rates that decrease toward the present.

### Historical biogeography

The model of biogeographic estimation with user-specified dispersal probabilities yielded a worse fit than the model with equal dispersal probabilities (likelihood with time-stratified dispersal multipliers: DEC_*strat*_=-143.41; likelihood without time-stratified dispersal multipliers DEC_*null*_=-140.75) and the ancestral state estimations involved some important differences. In both reconstructions the root state was highly unresolved. In the DEC_*null*_ model (highest likelihood), the area with the highest probability at the root was the southern part of the Amazonian Basin, *ca*. 28.1 Ma. The early divergence of the clade containing *M. marcus* and *M. eugenia* was accompanied by a colonization of the northern part of the Amazonian Basin (Fig. 4). The ancestor of the remaining group of *Morpho* occupied the Central Andes. This lineage then diverged into an Andean and an Amazonian lineage. This event (21.8 Ma) was also accompanied by a shift in microhabitat use: flight in low forest strata (understory) for the Andean lineage, and flight high above ground up to the canopy for the Amazonian lineage. The Andean lineage began a long-term occupation of the Central Andes with local diversification (12 nodes inferred occupying the Central Andes after the initial dispersal event). Around 11-12 Ma, cis-Andean (east of the Andes) recolonizations of Amazonia and the Atlantic Forest happened in three lineages. *M. polyphemus* is an intriguing case as it diverged 20.8 Ma from an Andean ancestor, but nowadays occupies Central America, whose connection to South America is often considered to be only completed during the last 4-3 million years. This implies either an earlier dispersal route of emerging Central America or a more recent dispersal with a joint extinction in the South American landmass. Overall, Northern Amazonia and the Northern Andes appear to have been colonized recently, during the last 5 million years (Fig. 4).

**Figure 4.**
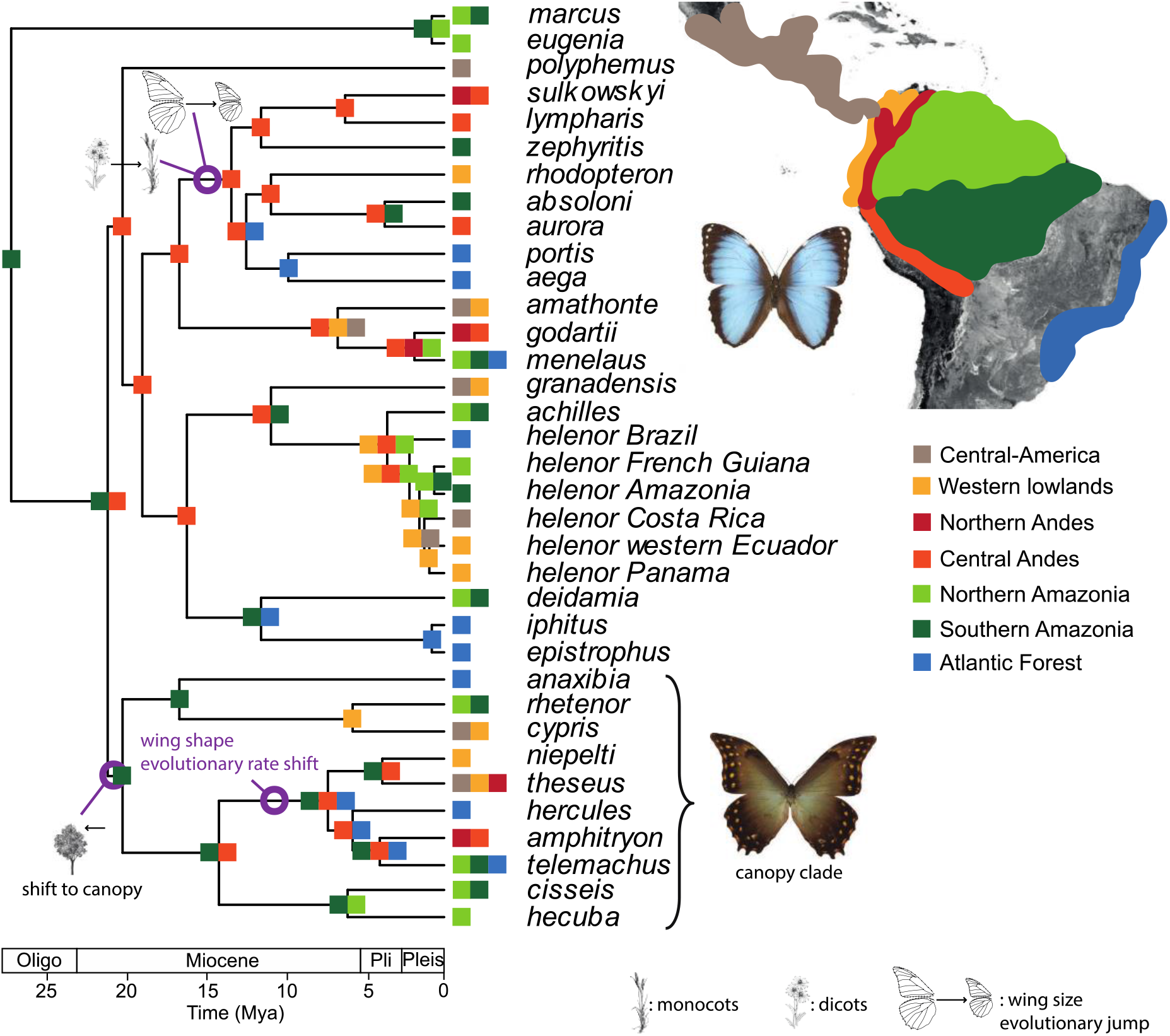
Historical biogeography inferred for the genus *Morpho*. The most likely states are indicated at the nodes. The different clade-specific ecological factors are also indicated on the tree. The two pictures of *Morpho* depict the typical wing shapes associated with each microhabitat – top: short rounded wings characteristic of the understory species, bottom: elongated wings toward the apex characteristic of the canopy clade.

## Discussion

In this study we aimed at investigating the large-scale patterns of diversification of the *Morpho* butterflies by jointly evaluating the dynamics of species and phenotypic diversification, to assess whether they are coupled or not and to test whether they correlate with clade-specific factors and/or biogeographic events. Our results show that ecological idiosyncrasies predominantly explain the pattern of diversification, instead of global (tree wide) factors. These ecological changes affected to a large extent both species and phenotypic diversification, leading to the partial coupling of both dynamics. Based on the amount of information currently available on the ecology of *Morpho* we discuss the potential role of several ecological and biogeographic events as well as the correlation with phenotypic diversification in explaining these variations among groups.

### Study limitations

A number of limitations have to be mentioned before discussing our results. Focusing on a small clade allowed us to combine multiple ecological, morphological and historical components thereby providing a deep understanding of the *Morpho* history. Although we sampled all known species for both the molecular phylogeny and morphological traits, our comparative analyses probably lack power as a result of both the small number of taxa (30 species) and the phylogenetic distribution of the traits of interest. Both microhabitat shift and host-plant shift (towards monocotyledons) are single events happening at the root of a single clade each and we lack phylogenetically independent similar shifts. Typically, we found an evolutionary jump in wing size to be associated with a shift from dicotyledons to monocotyledons host-plants. Further work addressing this pattern at a larger phylogenetic scale will be necessary to assess the generality of our finding. Furthermore, the reliability of birth-death models to assess the diversification dynamics from phylogenies of extant taxa is debated (e.g. Quental & Marshall, 2010; Louca & Pennell, 2020). We thus remain cautious with our estimation of the diversification dynamics and the interpretation of the different models tested. In particular, we avoided interpreting the speciation and extinction rates independently to focus only on the net diversification dynamics. Finally, the timing and magnitude of the Andean surface uplift is also controversial (see for example Evenstar *et al*.,2015, and references therein; Fiorella *et al*., 2015). We based our test on the reconstruction proposed by Leier *et al*. (2013) that focused only on the eastern cordillera of the Central Andes where the *Morpho* diversity is the highest, but had a large uncertainty in their paleo-altitude estimations. The Andean orogeny was spatially and temporally heterogeneous (Horton, 2018), which makes the use and interpretation of the paleoaltitude-dependent diversification model difficult (Condamine *et al*., 2018). Those limitations should thus be kept in mind throughout the following discussion of the drivers of diversification, and the signal of declining diversity in particular.

### Early Andean diversification not directly driven by Andean uplift

The diversification of the genus *Morpho* in the Andes could have happened either simultaneously with the uplift – a scenario where speciation is driven by the increasing heterogeneity of ecological conditions with new altitudes (Lagomarsino *et al*., 2016) – or decoupled from orogenesis – a scenario where a clade radiates across a range of altitudes already established through adaptations to ecological conditions (e.g. climate, host-plants, predators). Our results support the second hypothesis. We found that a model of diversification rate responding to paleo-altitude performed worse than the clade-specific diversification models (Tables 2 and 3), which means that neither global speciation nor extinction rate variations are well explained by the paleo-altitudes of the Central Andes. From a biogeographic point of view, 16 extant species (over 30) are almost restricted to the lowlands, while only six extant species have a distribution strictly restricted to the Andes. Yet, from the Oligocene-Miocene boundary to middle Miocene periods (23.5 to 11.6 Ma), 11 nodes out of 14 were inferred to be at least in the Central Andes from our biogeographic estimation (Fig. 4). Combined to the hypothesis that the *Morpho* probably originated in the foothills of the proto-Central-Andes, it is undeniable that the Central Andes played an important role in the early diversification of *Morpho*. During the second half of their evolutionary history, these lineages dispersed and diversified out of the Central Andes.

In contrast with the pattern of Central Andean diversification described above, the Northern Andes appear to have played only a minor role: while Northern Andean uplift likely established a barrier in three instances, resulting in cis- and trans-Andean *Morpho* lineages (Fig. 4), no major diversification was associated with the periods of Northern Andean uplift (Blandin & Purser, 2013). This absence of local diversification in the Northern Andes is a major difference compared to other butterflies such as the Ithomiini in which several groups repeatedly diversified at a high rate in the Northern Andes such as the genera *Napeogenes* (Elias *et al*., 2009), *Oleria* (De-Silva *et al*., 2016), *Hypomenitis* (Chazot *et al*., 2016) or *Pteronymia* (De-Silva *et al*., 2017).

Diversification driven by host-plant evolution may be an alternative explanation for the early diversification of *Morpho*. Penz and DeVries (2002) and Cassildé *et al*. (2010) suggested that monocotyledons were the ancestral host-plants of the genus *Morpho*, probably because at the time it was admitted that *M. marcus* larvae feed on monocotyledons (Constantino, 1997). However, we now know that *M. marcus* very probably feeds on Fabaceae (e.g. *Inga auristellae*; Ramírez-Garcia *et al*., 2014; Vásquez Bardales *et al*., 2017), and *M. eugenia* certainly feed on Caesalpiniaceae (Bénéluz, 2016). Therefore, since groups closely related to *Morpho*, notably the sister genus *Caerois*, are known to only feed on monocotyledon host-plants (Beccaloni *et al*., 2008), it is likely that the divergence of the *Morpho* was associated with an initial shift to dicotyledons. This host-plant shift at the root of Morphos created the conditions for an early rapid diversification of the group.

### A shift towards the canopy driving phenotypic and diversification changes

We found a shift of species diversification associated with a single shift from the understory to the canopy (DeVries *et al*. 2010; Chazot *et al*. 2016). We also found strong indications that female wing shape evolution in the canopy clade is different from a neutral evolution. An increasing rate of shape evolution for both fore- and hind-wings was supported in the subclade nested in the canopy clade and including *M. theseus*, *M. niepelti*, *M. amphytrion*, *M. telemachus*, and *M. hercules*. Chazot *et al*. (2016) showed that both male and female wing shapes in the canopy clade are significantly different from wing shapes in understory species. Here we show that this microhabitat change associated with different vegetation structure, microclimatic conditions and predator community may have also affected the rate of female wing shape evolution in addition to shape *per se*. However, we note that the highest rate shift was not placed at the root of the canopy clade, suggesting that other factors may have caused this rapid phenotypic evolution. This increased rate of wing shape evolution was not found in males. Instead, in males we found two significant slowdowns in rate at different small subclades, only in the case of hindwings. The lack of more precise information on these species ecology unfortunately prevents speculating on the factors involved in such changes in wing shape evolutionary rate.

### A second change in microhabitat conditions associated with a host-plant, phenotypical and diversification shifts

Published information in the *portis* clade (Heredia & Alvarez, 2007; Beccaloni *et al*., 2008; Montero Abril & Ortiz Perez, 2010) indicate that four *Morpho* species (*M. portis, M. aega, M. sulkowskyi* and *M. rhodopteron*) feed on Neotropical woody bamboos (Poaceae, tribe Bambuseae), notably on *Chusquea* species (subtribe Chusqueinae), in particular *Chusquea* aff. *scandens* for *M. sulkowskyi* that occurs at cloud forest elevations (Heredia & Alvarez, 2007). Recent observations indicate that M. zephyritis also feeds on woody bamboos (Roberto Maravi, pers. comm.). For the other species of the *portis* clade, there are only field observations indicating that they live in areas with important bamboo vegetation (Purser & Lacomme, 2016; pers. obs. in Peru, Daniel Lacomme pers. com.).

If, as observations indicate, the *portis* clade diversified after an initial shift back to monocotyledon host-plants, this reversal evolutionary event is a strong support for the “oscillation hypothesis” (Janz *et al*., 2006). This hypothesis was proposed to explain the pattern of nymphalid butterflies with respect to host-plant use (Janz *et al*., 2006) and states that the ability to recolonize “lost” hosts should be conserved over long evolutionary times, leading to recurrent recolonization events. Compared to the speciation rate of the backbone, species diversification within the *portis* clade proceeded at a higher rate, and rapidly decreased through time to reach almost zero at present. Adaptive radiations, here following a major host-plant shift, predict this rapid dampening of speciation rate as a result of niche filling (Schluter, 2000; Gavrilets & Losos, 2009).

Interestingly, we found that an evolutionary jump – a fast punctual event of evolution – toward smaller wing sizes also coincided with the host-plant shift. Chazot *et al*. (2016) did not identify any driver of this wing size evolution. To our knowledge, there is no clear expectation or evidence supporting a specific relationship between body size and monocot *versus* dicot feeders but this question has rarely been addressed (but see Garcia-Barros 2000). The jump toward smaller sizes also cannot be associated with any altitudinal change because some species of the clade only occur at low to mid altitudes (200-1500 m), while others occur at higher altitudes (1500-3500 m) (Blandin, 2007; Gayman *et al*., 2016).

Therefore, other hypotheses need to be explored, in particular that of a second possible change of microhabitat conditions. Many Bambusinae, in particular *Chusquea* species, form dense thickets, twigs and leaves creating inextricable tangles as a result from abundant vegetative branching at each node (Fisher, 2011; Fisher *et al*., 2014). Observational data on the behaviour of the bamboo feeding *Morpho* is scarce, but observations on *M. rhodopteron* (Montero Abril and Ortiz Perez, 2010; Purser and Lacomme, 2016), *M. sulkowskyi* (Heredia and Alvarez-Lopez 2007), and *M. aega* (Otero & Marigo, 1990) suggest that females are more often resting inside the *Chusquea* thickets while males are flying around (males, when resting, also stand in the vegetation). Moreover, Heredia & Alvarez-Lopez (2007) noted that *M. sulkowskyi* females having light and dark alternating stripes on wings ventral side are difficult to detect inside *Chusquea* thickets. More or less contrasted similar patterns exist in males and females of other species, except in *M. absoloni*. Therefore, we hypothesize that size reduction, associated to a more or less striped appearance of the ventral side, could be an adaptation to the microhabitat structure of dense woody bamboo thickets, highlighting once again the importance of the microhabitat conditions on species and trait evolution.

### Declining diversity in the Neotropical

Morpho When accounting for heterogeneity in diversification rates (isolating the two shifting subclades), the diversification dynamics for the remaining lineages was characterized by a negative net diversification rate, indicative of a declining diversity, mainly during the Miocene. Whether diversity decline can be accurately estimated only from phylogenies of extant species is a matter of debate (e.g. Quental & Marshall, 2010; but see Morlon *et al*.,2011). In the case of *Morpho*, this pattern may explain why some branches in the tree (such as the stem branch of *M. marcus* and *M. eugenia* or the branches leading to *M. anaxibia, M. deidamia*, or *M. polyphemus*) are surprisingly long. Extinct lineages may also explain why *M. polyphemus*, which diverged from its sister clade 20 Ma, is found in Central America, while colonization of Central America is often expected to be much more recent (but see Montes *et al*., 2015, Farris *et al*., 2011). Major landscape transformations during the Miocene in western Amazonia may explain this decline. Between 23-10 Ma, Western Amazonia transformed into a large wetland of lakes, swamps and shallow water, called the Pebas System (Wesselingh *et al*., 2001; Hoorn *et al*., 2010). The exact nature of the Pebas System is still under discussion but it was most likely unsuitable for terrestrial fauna (Salas-Gismondi *et al*.,2015). Evidence of extinction has been found from a west Amazonian fossil record, in particular with a major decrease of mammalian diversity at the transition between the Oligocene and the Miocene (Antoine *et al*., 2016), which is in line with the beginning of the diversity decline in *Morpho* (Fig. 3).

### Conclusion

Our results support a prevailing ecological basis for both species and phenotypic diversification in *Morpho* butterflies: (1) a major host-plant shift, which punctually affected wing size evolution and greatly affected species diversification dynamics (pattern of adaptive radiation), and (2) a microhabitat shift affecting species diversification and partially wing shape diversification. Therefore, to a large extent, the dynamics of species diversification and phenotypic diversification are coupled in *Morpho*, most likely as a result of two major ecological events. More importantly, we show that both species and phenotypic diversification in *Morpho* butterflies are better explained by multiple clade-specific factors instead of global abiotic drivers. Current methods for identifying drivers of diversification, based on model comparisons, are unable to test for potential interactions between drivers. Hence, our results do not exclude the possibility that the Andes played a role in diversification, but rather suggest that their effect on the shape of the phylogenetic tree was less significant than other factors. Nevertheless, the extent to which the effects of these ecological drivers can be generalised is unknown given the scale of our dataset. In particular future work at a larger phylogenetic scale should shed light on the importance of major host-plant transitions on the evolution of body size and the dynamics of diversification. Our study also highlights that both phenotypic and ecological information are of key relevance for understanding macroevolutionary patterns of diversification.

## Acknowledgments

The authors have no conflict of interest to declare. F.L.C. has benefited from an “Investissements d’Avenir” grant managed by Agence Nationale de la Recherche (CEBA, ref. ANR-10-LABX-25-01). P.B benefited from the Programme Pluriformation “Etat et structure phylogénétique de la biodiversité actuelle et fossile”.

