## Supporting Information for "Punctual ecological changes rather than global factors drive species diversification and the evolution of wing phenotypes in *Morpho* butterflies"

Supporting Information S1: Tree reconstructed and time-calibrated using BEAST. Node ages in red and the 95% HPD bars are indicated at the nodes. Grey numbers from 1 to 7 are the positions of the different time-constraints used. We used uniform distributions bounded by the 95% HPD inferred by Wahlberg et al. (2009). Upper and lower boundaries are indicated. In green are the node numbers corresponding to the tests for different wing shape evolutionary rates.

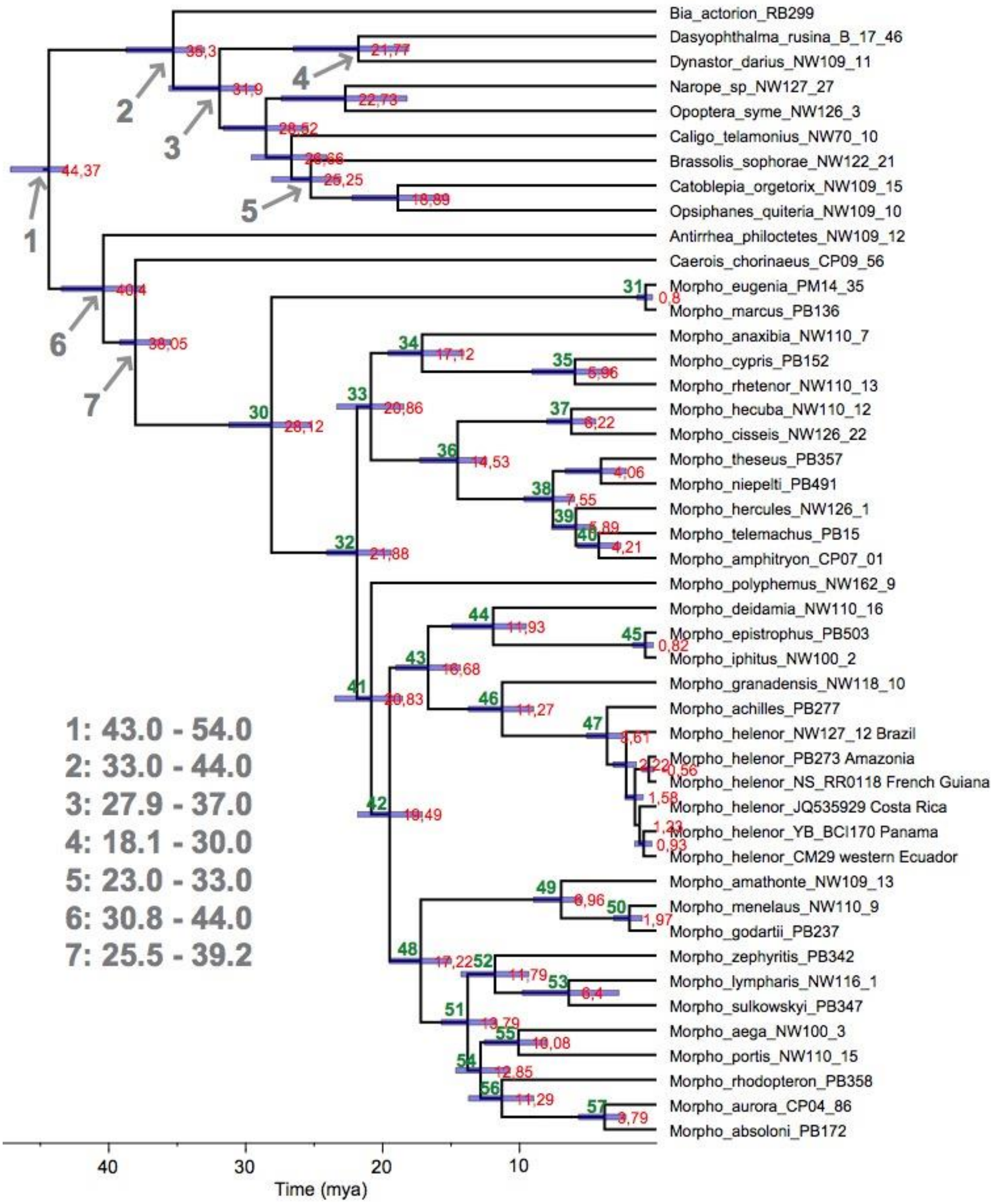

Supporting Information S2. Model comparison for wing size evolution for males and females for the analyses performed on the MCC tree.

a) Males

| | Models | AIC | ΔAIC |
|---|---|---|---|
| Forewing | bm | 126.833 | 0 |
| | rbm | 128.402 | 1.568 |
| | jbm | 153.859 | 27.025 |
| Hindwing | bm | 113.605 | 0 |
| | rbm | 115.105 | 1.5 |
| | jbm | 124.984 | 11.379 |

b) Females

| | Models | AIC | ΔAIC |
|---|---|---|---|
| Forewing | jbm | 122.68 | 0 |
| | bm | 154.069 | 31.389 |
| | rbm | 197.752 | 75.072 |
| Hindwing | jbm | 107.923 | 0 |
| | rbm | 117.123 | 9.2 |
| | bm | 197.547 | 89.624 |

Supporting Information S3. A. Results of models with shifts in rates of wing shape evolution for males on the MCC tree.

a) One-shift models

| **Wing** | **Node** | **Ingroup rate** | **Background rate** | **Ratio** | **p-value** |
|---|---|---|---|---|---|
| Forewing | 33 | 3.11E+17 | 3.70E+17 | 1.18 | 0.962 |
| | 34 | 5.01E+17 | 3.34E+17 | 1.50 | 0.927 |
| | 36 | 2.30E+17 | 3.87E+17 | 1.68 | 0.505 |
| | 38 | 2.05E+17 | 3.80E+17 | 1.85 | 0.228 |
| | 39 | 1.79E+17 | 3.70E+17 | 2.06 | 0.294 |
| | 42 | 2.92E+17 | 4.38E+17 | 1.49 | 0.452 |
| | 43 | 3.03E+17 | 4.13E+17 | 1.36 | 0.451 |
| | 44 | 2.62E+17 | 4.02E+17 | 1.53 | 0.846 |
| | 45 | 2.77E+17 | 3.77E+17 | 1.36 | 0.978 |
| | 46 | 3.19E+17 | 3.54E+17 | 1.11 | 0.985 |
| | 48 | 2.52E+17 | 3.70E+17 | 1.47 | 0.918 |
| | 49 | 2.93E+17 | 3.57E+17 | 1.21 | 0.933 |
| | 52 | 2.22E+17 | 3.65E+17 | 1.64 | 0.332 |
| | 54 | 3.78E+17 | 3.44E+17 | 1.09 | 0.912 |
| | 55 | 2.69E+17 | 3.60E+17 | 1.33 | 0.414 |
| | 57 | 4.87E+17 | 3.35E+17 | 1.45 | 0.672 |
| Hindwing | 33 | 5.6627E+17 | 5.4632E+17 | 1.03 | 0.994 |
| | 34 | 7.0283E+17 | 5.3632E+17 | 1.31 | 0.974 |
| | 36 | 5.0774E+17 | 5.6674E+17 | 1.11 | 0.93 |
| | 38 | 5.5832E+17 | 5.519E+17 | 1.01 | 0.984 |
| | 39 | 4.2922E+17 | 5.6672E+17 | 1.32 | 0.751 |
| | 42 | 4.7833E+17 | 6.6492E+17 | 1.39 | 0.536 |
| | 43 | 4.9549E+17 | 6.2814E+17 | 1.26 | 0.577 |
| | 44 | 5.9163E+17 | 5.3059E+17 | 1.11 | 0.991 |
| | 45 | 7.2644E+17 | 4.8989E+17 | 1.48 | 0.965 |
| | 46 | 8.6181E+17 | 5.1865E+17 | 1.66 | 0.852 |
| | 48 | 6.4522E+17 | 5.3452E+17 | 1.20 | 0.987 |
| | 49 | 7.0738E+17 | 5.3581E+17 | 1.32 | 0.892 |
| | 52 | 2.3213E+17 | 5.8862E+17 | 2.53 | 0.059 |
| | 54 | 3.1922E+17 | 6.1141E+17 | 1.91 | 0.229 |
| | **55** | **2.522E+17** | **5.8639E+17** | **2.32** | **0.02** |
| | 57 | 3.8624E+17 | 5.7149E+17 | 1.47 | 0.658 |

*Node = node tested for a shift (see Supporting Information S1). Ingroup rate = rate of evolution estimated in the putative shifting clade. Background rate = rate of evolution for the remaining phylogeny outside of the putative shifting clade. Ratio = ratio between the ingroup and the background rate. p-value = significance value of the ratio. Bold p-values indicate cases where the p-value was below the significance threshold of 0.05.*

b) Two-shifts models - hindwing

| Model | Structure | Rate | Pairwise comparison | Pairwise ratio | Pairwise p-value |
|---|---|---|---|---|---|
| | 55 | 2.52E+17 | 1_2 | 2.25 | 0.29 |
| 1 | 33 | 5.66E+17 | 1_B | 2.37 | 0.02 |
| | background | 5.98E+17 | 2_B | 1.06 | 0.99 |
| | 55 | 2.52E+17 | 1_2 | 2.79 | 0.44 |
| 2 | 34 | 7.02E+17 | 1_B | 2.27 | 0.02 |
| | background | 5.71E+17 | 2_B | 1.23 | 0.97 |
| | 55 | 2.52E+17 | 1_2 | 2.01 | 0.29 |
| 3 | 36 | 5.07E+17 | 1_B | 2.43 | 0.02 |
| | background | 6.13E+17 | 2_B | 1.21 | 0.88 |
| | 55 | 2.52E+17 | 1_2 | 2.21 | 0.15 |
| 4 | 38 | 5.58E+17 | 1_B | 2.35 | 0.02 |
| | background | 5.92E+17 | 2_B | 1.06 | 0.92 |
| | 55 | 2.52E+17 | 1_2 | 1.70 | 0.48 |
| 5 | 39 | 4.29E+17 | 1_B | 2.40 | 0.02 |
| | background | 6.06E+17 | 2_B | 1.41 | 0.67 |
| | 55 | 2.52E+17 | 1_2 | 2.35 | 0.26 |
| 6 | 44 | 5.91E+17 | 1_B | 2.31 | 0.03 |
| | background | 5.82E+17 | 2_B | 1.02 | 1.00 |
| | 55 | 2.52E+17 | 1_2 | 2.88 | 0.24 |
| 7 | 45 | 7.26E+17 | 1_B | 2.09 | 0.06 |
| | background | 5.27E+17 | 2_B | 1.38 | 0.99 |
| | 55 | 2.52E+17 | 1_2 | 3.42 | 0.22 |
| 8 | 46 | 8.61E+17 | 1_B | 2.19 | 0.02 |
| | background | 5.51E+17 | 2_B | 1.56 | 0.86 |
| | 55 | 2.52E+17 | 1_2 | 2.56 | 0.39 |
| 9 | 48 | 6.45E+17 | 1_B | 2.27 | 0.01 |
| | background | 5.73E+17 | 2_B | 1.13 | 0.99 |
| | 55 | 2.52E+17 | 1_2 | 2.80 | 0.23 |
| 10 | 49 | 7.07E+17 | 1_B | 2.27 | 0.03 |
| | background | 5.71E+17 | 2_B | 1.24 | 0.91 |
| | 55 | 2.52E+17 | 1_2 | 1.09 | 0.89 |
| 11 | 52 | 2.32E+17 | **1_B** | **2.50** | **0.01** |
| | background | 6.30E+17 | **2_B** | **2.72** | **0.04** |
| | 55 | 2.52E+17 | 1_2 | 1.53 | 0.57 |
| 12 | 57 | 3.86E+17 | 1_B | 2.42 | 0.02 |
| | background | 6.11E+17 | 2_B | 1.58 | 0.62 |

*Structure = structure of the model, i.e. the node number of the two putative shifts tested and remaining phylogeny outside from the putative shifting clade ("background"). In each case the first node indicated is the node retained in the one-shift model comparisons. Rate = rate of evolution of each part of the model, i.e. two ingroup rates and one background rate. Pairwise comparison = pairs considered in the following pairwise comparison (1-2=ingroup 1 and ingroup 2, 1-B=ingroup 1 and background, 2-B=ingroup 2 and background). Pairwise ratios = ratio between the two rates considered in the pairwise comparison. Pairwise p-value = significance value of the pairwise ratio. Bold p-values indicate the case for which both subclades were significantly different than the background rate.*

Supporting Information S4.

A. Results of models with shifts in rates of wing shape evolution for females on the MCC tree.

a) One-shift models

| Wing | Node | Ingroup rate | Background rate | Ratio | p-value |
|---|---|---|---|---|---|
| Forewing | 32 | 5.49E+19 | 5.40E+17 | 101.55 | 0.001 |
| | 33 | 5.58E+17 | 1.93E+19 | 34.67 | 0.001 |
| | 35 | 8.20E+19 | 5.43E+17 | 151.17 | 0.001 |
| | **37** | **1.22E+20** | **6.18E+17** | **197.98** | **0.001** |
| | 38 | 1.60E+20 | 9.45E+17 | 169.38 | 0.001 |
| | 40 | 5.18E+17 | 4.50E+19 | 86.97 | 0.001 |
| | 41 | 5.04E+17 | 4.13E+19 | 82.09 | 0.001 |
| | 42 | 6.09E+17 | 2.77E+19 | 45.45 | 0.001 |
| | 43 | 5.96E+17 | 2.38E+19 | 39.97 | 0.001 |
| | 44 | 6.17E+17 | 1.93E+19 | 31.35 | 0.001 |
| | 46 | 5.83E+17 | 2.09E+19 | 35.87 | 0.001 |
| | 47 | 5.78E+17 | 1.93E+19 | 33.48 | 0.001 |
| | 50 | 6.44E+17 | 1.93E+19 | 30.03 | 0.001 |
| | 52 | 3.11E+17 | 2.19E+19 | 70.32 | 0.001 |
| | 55 | 2.84E+17 | 1.94E+19 | 68.29 | 0.001 |
| Hindwing | 32 | 8.32E+19 | 8.50E+17 | 97.89 | 0.001 |
| | 33 | 7.17E+17 | 2.93E+19 | 40.99 | 0.001 |
| | 35 | 1.25E+20 | 8.33E+17 | 149.48 | 0.001 |
| | **37** | **1.86E+20** | **9.29E+17** | **199.94** | **0.001** |
| | 38 | 2.43E+20 | 1.47E+18 | 164.93 | 0.001 |
| | 40 | 8.57E+17 | 6.82E+19 | 79.66 | 0.001 |
| | 41 | 8.56E+17 | 6.26E+19 | 73.13 | 0.001 |
| | 42 | 9.60E+17 | 4.19E+19 | 43.71 | 0.001 |
| | 43 | 1.00E+18 | 3.61E+19 | 36.09 | 0.001 |
| | 44 | 9.08E+17 | 2.93E+19 | 32.31 | 0.001 |
| | 46 | 1.05E+18 | 3.17E+19 | 30.04 | 0.001 |
| | 47 | 1.01E+18 | 2.93E+19 | 28.79 | 0.001 |
| | 50 | 8.54E+17 | 2.93E+19 | 34.39 | 0.001 |
| | 52 | 6.66E+17 | 3.32E+19 | 49.75 | 0.001 |
| | 55 | 8.44E+17 | 2.93E+19 | 34.79 | 0.001 |

*Structure = structure of the model, i.e. the node number of the two putative shifts tested and remaining phylogeny outside from the putative shifting clade ("background"). In each case the first node indicated is the node retained in the one-shift model comparisons. Rate = rate of evolution of each part of the model, i.e. two subclade rates and one background rate. Pairwise comparison = pairs considered in the following pairwise comparison (1_2 = subclade 1 and subclade 2, 1_B = subclade 1 and background, 2_B = subclade 2 and background). Pairwise ratios = ratio between the two rates considered in the pairwise comparison. Pairwise p-value = significance value of the pairwise ratio. Bold p-values indicate the case for which both subclades were significantly different than the background rate.*

b) Two-shifts models - forewing

| Model | Structure | Rate | Pairwise comparison | Pairwise ratio | Pairwise p-value |
|---|---|---|---|---|---|
| | 37 | 1.22E+20 | 1_2 | 219.22 | 0.001 |
| 1 | 33 | 5.57E+17 | 1_B | 195.40 | 0.001 |
| | background | 6.25E+17 | 2_B | 1.12 | 0.95 |
| | 37 | 1.22E+20 | 1_2 | 236.21 | 0.001 |
| 2 | 40 | 5.17E+17 | 1_B | 139.80 | 0.001 |
| | background | 8.74E+17 | 2_B | 1.68 | 0.521 |
| | 37 | 1.22E+20 | 1_2 | 242.87 | 0.001 |
| 3 | 41 | 5.03E+17 | 1_B | 142.15 | 0.001 |
| | background | 8.60E+17 | 2_B | 1.70 | 0.469 |
| | 37 | 1.22E+20 | 1_2 | 200.93 | 0.001 |
| 4 | 42 | 6.08E+17 | 1_B | 195.72 | 0.001 |
| | background | 6.24E+17 | 2_B | 1.02 | 0.99 |
| | 37 | 1.22E+20 | 1_2 | 205.38 | 0.001 |
| 5 | 43 | 5.95E+17 | 1_B | 194.68 | 0.001 |
| | background | 6.28E+17 | 2_B | 1.05 | 0.983 |
| | 37 | 1.22E+20 | 1_2 | 198.28 | 0.001 |
| 6 | 44 | 6.16E+17 | 1_B | 197.94 | 0.001 |
| | background | 6.17E+17 | 2_B | 1.00 | 1 |
| | 37 | 1.22E+20 | 1_2 | 209.89 | 0.001 |
| 7 | 46 | 5.82E+17 | 1_B | 195.21 | 0.001 |
| | background | 6.26E+17 | 2_B | 1.07 | 0.964 |
| | 37 | 1.22E+20 | 1_2 | 211.71 | 0.001 |
| 8 | 47 | 5.77E+17 | 1_B | 196.25 | 0.001 |
| | background | 6.23E+17 | 2_B | 1.07 | 0.942 |
| | 37 | 1.22E+20 | 1_2 | 189.97 | 0.001 |
| 9 | 50 | 6.43E+17 | 1_B | 199.13 | 0.001 |
| | background | 6.14E+17 | 2_B | 1.04 | 0.96 |
| | 37 | 1.22E+20 | 1_2 | 393.42 | 0.001 |
| 10 | 52 | 3.10E+17 | 1_B | 171.13 | 0.001 |
| | background | 7.14E+17 | 2_B | 2.29 | 0.231 |
| | 37 | 1.22E+20 | 1_2 | 361.82 | 0.001 |
| 11 | 53 | 3.38E+17 | 1_B | 186.47 | 0.001 |
| | background | 6.55E+17 | 2_B | 1.94 | 0.328 |
| | 37 | 1.22E+20 | 1_2 | 431.07 | 0.001 |
| 12 | 55 | 2.83E+17 | 1_B | 184.39 | 0.001 |
| | background | 6.63E+17 | 2_B | 2.33 | 0.333 |

*Structure = structure of the model, i.e. the node number of the two putative shifts tested and remaining phylogeny outside from the putative shifting clade ("background"). In each case the first node indicated is the node retained in the one-shift model comparisons. Rate = rate of evolution of each part of the model, i.e. two subclade rates and one background rate. Pairwise comparison = pairs considered in the following pairwise comparison (1_2 = subclade 1 and subclade 2, 1_B = subclade 1 and background, 2_B = subclade 2 and background). Pairwise ratios = ratio between the two rates considered in the pairwise comparison. Pairwise p-value = significance value of the pairwise ratio. Bold p-values indicate the case for which both subclades were significantly different than the background rate.*

c) Two-shifts models - hindwing

| Model | Structure | Rate | Pairwise comparison | Pairwise ratio | Pairwise p-value |
|---|---|---|---|---|---|
| | 37 | 1.86E+20 | 1_2 | 259.14 | 0.001 |
| 1 | 33 | 7.17E+17 | 1_B | 193.90 | 0.001 |
| | background | 9.58E+17 | 2_B | 1.33 | 0.892 |
| | 37 | 1.86E+20 | 1_2 | 216.81 | 0.001 |
| 2 | 40 | 8.57E+17 | 1_B | 166.60 | 0.001 |
| | background | 1.11E+18 | 2_B | 1.30 | 0.747 |
| | 37 | 1.86E+20 | 1_2 | 216.87 | 0.001 |
| 3 | 41 | 8.56E+17 | 1_B | 171.49 | 0.001 |
| | background | 1.08E+18 | 2_B | 1.26 | 0.733 |
| | 37 | 1.86E+20 | 1_2 | 193.41 | 0.001 |
| 4 | 42 | 9.60E+17 | 1_B | 205.38 | 0.001 |
| | background | 9.04E+17 | 2_B | 1.06 | 0.977 |
| | 37 | 1.86E+20 | 1_2 | 185.70 | 0.001 |
| 5 | 43 | 1.00E+18 | 1_B | 207.42 | 0.001 |
| | background | 8.95E+17 | 2_B | 1.11 | 0.964 |
| | 37 | 1.86E+20 | 1_2 | 204.42 | 0.001 |
| 6 | 44 | 9.08E+17 | 1_B | 199.34 | 0.001 |
| | background | 9.32E+17 | 2_B | 1.02 | 0.988 |
| | 37 | 1.86E+20 | 1_2 | 176.03 | 0.001 |
| 7 | 46 | 1.05E+18 | 1_B | 206.96 | 0.001 |
| | background | 8.97E+17 | 2_B | 1.17 | 0.928 |
| | 37 | 1.86E+20 | 1_2 | 182.23 | 0.001 |
| 8 | 47 | 1.01E+18 | 1_B | 202.62 | 0.001 |
| | background | 9.17E+17 | 2_B | 1.11 | 0.946 |
| | 37 | 1.86E+20 | 1_2 | 217.50 | 0.001 |
| 9 | 50 | 8.54E+17 | 1_B | 197.76 | 0.001 |
| | background | 9.39E+17 | 2_B | 1.09 | 0.935 |
| | 37 | 1.86E+20 | 1_2 | 278.86 | 0.001 |
| 10 | 52 | 6.66E+17 | 1_B | 183.53 | 0.001 |
| | background | 1.01E+18 | 2_B | 1.51 | 0.536 |
| | 37 | 1.86E+20 | 1_2 | 380.70 | 0.001 |
| 11 | 53 | 4.88E+17 | 1_B | 187.78 | 0.001 |
| | background | 9.89E+17 | 2_B | 2.02 | 0.336 |
| | 37 | 1.86E+20 | 1_2 | 220.01 | 0.001 |
| 12 | 55 | 8.44E+17 | 1_B | 197.48 | 0.001 |
| | background | 9.40E+17 | 2_B | 1.11 | 0.882 |

*Structure = structure of the model, i.e. the node number of the two putative shifts tested and remaining phylogeny outside from the putative shifting clade (“background”). In each case the first node indicated is the node retained in the one-shift model comparisons. Rate = rate of evolution of each part of the model, i.e. two subclade rates and one background rate. Pairwise comparison = pairs considered in the following pairwise comparison (1_2 = subclade 1 and subclade 2, 1_B = subclade 1 and background, 2_B = subclade 2 and background). Pairwise ratios = ratio between the two rates considered in the pairwise comparison. Pairwise p-value = significance value of the pairwise ratio. Bold p-values indicate the case for which both subclades were significantly different than the background rate.*

Supporting Information S5. Results of all time-dependent diversification models fitted on the *Morpho*, the different subscales tested (canopy, monocots, shape shift) and the corresponding backbone trees. *Model* indicates the shape of speciation and extinction rate functions: BCST=constant speciation, BVAR=time-dependent speciation, DCST=constant extinction, DVAR=time-dependent extinction. Par=number of parameters in the model. logL=likelihood of the model. AIC=Akaike Information Criterion. λ =speciation rate at present, α=coefficient of time variation of the speciation rate, µ =extinction rate at present, ß =coefficient of time variation of the extinction rate.

**Whole tree**

| Model | Par | logL | AIC | ΔAIC | λ | α | µ | β |
|---|---|---|---|---|---|---|---|---|
| BVAR | 2 | -96.97 | 197.95 | 0.00 | 0.053 | 0.047 | | |
| BCST | 1 | -98.40 | 198.81 | 0.86 | 0.081 | | | |
| BVARDCST | 3 | -96.97 | 199.95 | 2.00 | 0.053 | 0.047 | 0.000 | |
| BCSTDCST | 2 | -98.40 | 200.81 | 2.86 | 0.081 | | 0.000 | |
| BVARDVAR | 4 | -96.97 | 201.95 | 4.00 | 0.053 | 0.047 | 0.000 | 0.019 |
| BCSTDVAR | 3 | -98.40 | 202.81 | 4.86 | 0.081 | | 0.000 | 0.014 |

**Background without canopy**

| Model | Par | logL | AIC | ΔAIC | λ | α | µ | β |
|---|---|---|---|---|---|---|---|---|
| BCST | 1 | -67.01 | 136.03 | 0.00 | 0.080 | | | |
| BVAR | 2 | -66.10 | 136.20 | 0.18 | 0.053 | 0.044 | | |
| BCSTDCST | 2 | -67.01 | 138.03 | 2.00 | 0.080 | | 0.000 | |
| BVARDCST | 3 | -66.04 | 138.09 | 2.06 | 0.060 | 0.049 | 0.034 | |
| BVARDVAR | 4 | -65.75 | 139.50 | 3.47 | 0.144 | 0.006 | 1.048 | -0.691 |
| BCSTDVAR | 3 | -67.01 | 140.03 | 4.00 | 0.080 | | 0.000 | 0.002 |

**Background without monocots**

| Model | Par | logL | AIC | ΔAIC | λ | α | µ | β |
|---|---|---|---|---|---|---|---|---|
| BCST | 1 | -73.75 | 149.49 | 0.00 | 0.081 | | | |
| BVAR | 2 | -73.24 | 150.48 | 0.98 | 0.060 | 0.031 | | |
| BVARDVAR | 4 | -71.58 | 151.16 | 1.66 | 0.034 | 0.205 | 0.039 | 0.201 |
| BCSTDCST | 2 | -73.75 | 151.49 | 2.00 | 0.081 | | 0.000 | |
| BVARDCST | 3 | -72.83 | 151.65 | 2.16 | 0.081 | 0.051 | 0.101 | |
| BCSTDVAR | 3 | -73.75 | 153.49 | 4.00 | 0.081 | | 0.000 | -0.039 |

**Background without shape shift**

| Model | Par | logL | AIC | ΔAIC | λ | α | µ | β |
|---|---|---|---|---|---|---|---|---|
| BVAR | 2 | -76.57 | 157.13 | 0.00 | 0.045 | 0.054 | | |
| BCST | 1 | -78.18 | 158.36 | 1.22 | 0.078 | | | |
| BVARDCST | 3 | -76.52 | 159.04 | 1.91 | 0.051 | 0.059 | 0.028 | |

| | | | | | | | | |
|---|---|---|---|---|---|---|---|---|
| BCSTDCST | 2 | -78.18 | 160.36 | 3.22 | 0.078 | | 0.000 | |
| BVARDVAR | 4 | -76.45 | 160.89 | 3.76 | 0.061 | 0.047 | 0.146 | -0.287 |
| BCSTDVAR | 3 | -78.18 | 162.36 | 5.22 | 0.078 | | 0.000 | -0.008 |

**Background without monocots and canopy**

| Model | Par | logL | AIC | ΔAIC | λ | α | μ | β |
|---|---|---|---|---|---|---|---|---|
| BVARDVAR | 4 | -35.68 | 79.36 | 0.00 | 0.063 | 0.237 | 0.079 | 0.228 |
| BCST | 1 | -39.78 | 81.57 | 2.20 | 0.073 | | | |
| BCSTDCST | 2 | -39.39 | 82.78 | 3.41 | 0.119 | | 0.089 | |
| BCSTDVAR | 3 | -38.70 | 83.39 | 4.03 | 0.180 | | 0.493 | -0.154 |
| BVARDCST | 3 | -39.47 | 84.94 | 5.57 | 0.183 | 0.045 | 0.330 | |
| BVAR | 2 | -42.22 | 88.43 | 9.07 | 0.064 | 0.020 | | |

**Background without monocots and shape shift**

| Model | Par | logL | AIC | ΔAIC | λ | α | μ | β |
|---|---|---|---|---|---|---|---|---|
| BCST | 1 | -50.91 | 103.83 | 0.00 | 0.072 | | | |
| BVARDVAR | 4 | -48.05 | 104.11 | 0.28 | 0.046 | 0.215 | 0.061 | 0.205 |
| BCSTDVAR | 3 | -49.56 | 105.13 | 1.30 | 0.177 | | 0.668 | -0.242 |
| BCSTDCST | 2 | -50.90 | 105.80 | 1.97 | 0.078 | | 0.013 | |
| BVARDCST | 3 | -50.72 | 107.45 | 3.62 | 0.124 | 0.057 | 0.246 | |
| BVAR | 2 | -52.90 | 109.80 | 5.97 | 0.050 | 0.039 | | |

**Canopy**

| Model | Par | logL | AIC | ΔAIC | λ | α | μ | β |
|---|---|---|---|---|---|---|---|---|
| BCST | 1 | -31.39 | 64.77 | 0.00 | 0.083 | | | |
| BVAR | 2 | -30.81 | 65.62 | 0.84 | 0.050 | 0.059 | | |
| BCSTDCST | 2 | -31.39 | 66.77 | 2.00 | 0.083 | | 0.000 | |
| BVARDCST | 3 | -30.81 | 67.62 | 2.84 | 0.050 | 0.059 | 0.000 | |
| BCSTDVAR | 3 | -31.39 | 68.77 | 4.00 | 0.083 | | 0.000 | 0.014 |
| BVARDVAR | 4 | -30.81 | 69.62 | 4.84 | 0.050 | 0.059 | 0.000 | 0.020 |

**Monocots**

| Model | Par | logL | AIC | ΔAIC | λ | α | μ | β |
|---|---|---|---|---|---|---|---|---|
| BVAR | 2 | -21.78 | 47.55 | 0.00 | 0.014 | 0.213 | | |
| BVARDCST | 3 | -21.78 | 49.55 | 2.00 | 0.014 | 0.213 | 0.000 | |
| BCST | 1 | -24.66 | 51.31 | 3.76 | 0.080 | | | |
| BVARDVAR | 4 | -21.78 | 51.55 | 4.00 | 0.014 | 0.213 | 0.000 | 0.010 |
| BCSTDCST | 2 | -24.66 | 53.31 | 5.76 | 0.080 | | 0.000 | |
| BCSTDVAR | 3 | -24.66 | 55.31 | 7.76 | 0.080 | | 0.000 | 0.010 |

**Shape shift**

| Model | Par | logL | AIC | ΔAIC | λ | α | μ | β |
|---|---|---|---|---|---|---|---|---|
| BCST | 1 | -20.14 | 42.28 | 0.00 | 0.095 | | | |

| | | | | | | | | |
|---|---|---|---|---|---|---|---|---|
| BVAR | 2 | -20.09 | 44.18 | 1.91 | 0.081 | 0.023 | | |
| BCSTDCST | 2 | -20.14 | 44.28 | 2.00 | 0.095 | | 0.000 | |
| BVARDCST | 3 | -20.09 | 46.18 | 3.91 | 0.081 | 0.023 | 0.000 | |
| BCSTDVAR | 3 | -20.14 | 46.28 | 4.00 | 0.095 | | 0.000 | 0.029 |
| BVARDVAR | 4 | -20.09 | 48.18 | 5.91 | 0.081 | 0.023 | 0.000 | 0.037 |

Supporting Information S6. Adjacency matrix and dispersal matrices time-stratified used for the BioGeoBEARS ancestral state reconstruction. CA=Central America, SN=Sierra Nevada de Santa Maria, W=lowland western Andes, N=Northern Andes, S=Central Andes, NAz=Northern Amazonia, SAZ=Southern Amazonia, AF=Atlantic Forest.

a) Adjacency matrix

| | CA | SN | W | N | S | NAz | SAz | AF |
|---|---|---|---|---|---|---|---|---|
| CA | 1 | 0 | 1 | 1 | 0 | 0 | 0 | 0 |
| SN | 0 | 1 | 1 | 0 | 0 | 0 | 0 | 0 |
| W | 1 | 1 | 1 | 1 | 1 | 1 | 0 | 0 |
| N | 1 | 1 | 1 | 1 | 1 | 1 | 0 | 0 |
| S | 0 | 1 | 1 | 1 | 0 | 0 | 1 | 1 |
| NAz | 0 | 1 | 1 | 0 | 1 | 1 | 1 | 0 |
| SAz | 0 | 0 | 0 | 1 | 1 | 1 | 1 | 1 |
| AF | 0 | 0 | 0 | 1 | 0 | 0 | 1 | 1 |

b) Dispersal multipliers matrix

**0-5 mya**

| | CA | SN | W | N | S | NAz | SAz | AF |
|---|---|---|---|---|---|---|---|---|
| CA | 1 | 0 | 1 | 0 | 0 | 0 | 0 | 0 |
| SN | 0 | 1 | 0 | 0 | 0 | 0 | 0 | 0 |
| W | 1 | 0 | 1 | 0 | 0 | 0 | 0 | 0 |
| N | 0 | 0 | 0 | 1 | 1 | 1 | 1 | 0 |
| S | 0 | 0 | 0 | 1 | 1 | 0.1 | 1 | 0.5 |
| NAz | 0 | 0 | 0 | 1 | 0.1 | 1 | 0.1 | 0 |
| SAz | 0 | 0 | 0 | 1 | 1 | 0.1 | 1 | 0 |
| AF | 0 | 0 | 0 | 0 | 0.5 | 0 | 0 | 1 |

**5-10 mya**

| | CA | SN | W | N | S | NAz | SAz | AF |
|---|---|---|---|---|---|---|---|---|
| CA | 1 | 0 | 0 | 0 | 0 | 0 | 0 | 0 |
| SN | 0 | 1 | 0 | 0 | 0 | 0 | 0 | 0 |
| W | 0 | 0 | 1 | 0.5 | 0.5 | 0.25 | 0 | 0 |
| N | 0 | 0 | 0.5 | 1 | 0.5 | 0.5 | 0.5 | 0 |
| S | 0 | 0 | 0.5 | 0.5 | 1 | 0.1 | 1 | 0.5 |
| NAz | 0 | 0 | 0.25 | 0.5 | 0.1 | 1 | 0.1 | 0 |
| SAz | 0 | 0 | 0 | 0.5 | 1 | 0.1 | 1 | 0 |
| AF | 0 | 0 | 0 | 0 | 0.5 | 0 | 0 | 1 |

**10-15 mya**

| | CA | SN | W | N | S | NAz | SAz | AF |
|---|---|---|---|---|---|---|---|---|
| CA | 1 | 0 | 0 | 0 | 0 | 0 | 0 | 0 |
| SN | 0 | 1 | 0.5 | 0.25 | 0 | 0 | 0 | 0 |
| W | 0 | 0.5 | 1 | 0.5 | 0.5 | 0 | 0 | 0 |
| N | 0 | 0.25 | 0.5 | 1 | 0 | 0 | 0 | 0 |
| S | 0 | 0 | 0.5 | 0 | 1 | 1 | 1 | 0.5 |
| NAz | 0 | 0 | 0 | 0 | 1 | 1 | 1 | 0 |
| SAz | 0 | 0 | 0 | 0 | 1 | 1 | 1 | 0 |
| AF | 0 | 0 | 0 | 0 | 0.5 | 0 | 0 | 1 |

**15-23 mya**

| | CA | SN | W | N | S | NAz | SAz | AF |
|---|---|---|---|---|---|---|---|---|
| CA | 1 | 0 | 0.5 | 0 | 0 | 0 | 0 | 0 |
| SN | 0 | 1 | 0.5 | 0.5 | 0 | 0 | 0 | 0 |
| W | 0.5 | 0.5 | 1 | 1 | 0.5 | 0 | 0 | 0 |
| N | 0 | 0.5 | 1 | 1 | 0 | 0 | 0 | 0 |
| S | 0 | 0 | 0.5 | 0 | 1 | 1 | 1 | 0.5 |
| NAz | 0 | 0 | 0 | 0 | 1 | 1 | 1 | 0 |
| SAz | 0 | 0 | 0 | 0 | 1 | 1 | 1 | 0 |
| AF | 0 | 0 | 0 | 0 | 0.5 | 0 | 0 | 1 |

**23-30 mya**

| | CA | SN | W | N | S | NAz | SAz | AF |
|---|---|---|---|---|---|---|---|---|
| CA | 1 | 0 | 0 | 0 | 0 | 0 | 0 | 0 |
| SN | 0 | 1 | 0.5 | 0.5 | 0 | 0 | 0 | 0 |
| W | 0 | 0.5 | 1 | 1 | 0.5 | 0 | 0 | 0 |
| N | 0 | 0.5 | 1 | 1 | 0.5 | 0 | 0.5 | 0 |
| S | 0 | 0 | 0.5 | 0.5 | 1 | 1 | 1 | 0.5 |
| NAz | 0 | 0 | 0 | 0 | 1 | 1 | 1 | 0.5 |
| SAz | 0 | 0 | 0 | 0.5 | 1 | 1 | 1 | 0.5 |
| AF | 0 | 0 | 0 | 0 | 0.5 | 0.5 | 0.5 | 1 |
